# Fourth generation CAR Tregs with *PDCD1*-driven IL-10 have enhanced suppressive function

**DOI:** 10.1101/2024.10.01.616177

**Authors:** Dominic A Boardman, Sonya Mangat, Jana K Gillies, Vivian CW Fung, Manjurul Haque, Majid Mojibian, Karoliina Tuomela, Christine M Wardell, Andrew Brown, Avery J Lam, Megan K Levings

## Abstract

The potency of regulatory T cell (Treg) therapy has been transformed through use of chimeric antigen receptors (CAR). However, to date, CAR Treg therapy has not achieved long-lasting tolerance in mouse models, suggesting that additional engineering is required to unlock the full potential of these cells. We previously found that human Tregs produce minimal amounts of IL-10 and have a limited capacity to control innate immunity in comparison to type I regulatory (Tr1) cells. Seeking to create “hybrid” CAR Tregs that were engineered with Tr1-like properties, we examined whether the *PDCD1* locus could be exploited to endow Tregs with the ability to secrete high levels of IL-10 in a CAR-regulated manner. CRISPR-mediated PD1-deletion increased the activation potential of CAR Tregs without compromising *in vivo* stability. Knock-in of *IL10* under control of the PD1 promoter facilitated CAR-mediated secretion of IL-10 in large quantities, and improved CAR Treg function, as determined by significant inhibition of dendritic cell antigen presentation and enhanced suppression of alloantigen- and islet autoantigen-specific T cells. Overall, CRISPR-mediated engineering to simultaneously remove an inhibitory signal and enhance suppressive mechanisms is a new approach to enhance the therapeutic potency of CAR Tregs.

## INTRODUCTION

Adoptive transfer of regulatory T cells (Treg) is a promising therapeutic strategy for treating transplant rejection, as well as a variety of autoimmune diseases and inflammatory disorders. Clinical trials have shown that polyclonal Treg therapy is feasible and well-tolerated (*1, 2*) but it is generally thought that efficacy is limited without enrichment for antigen-specific cells. Numerous groups have thus explored the potential of using chimeric antigen receptors (CAR) to confer disease-relevant antigen specificity to Tregs. CARs are synthetic fusion proteins that use domains from various receptors to redirect the specificity of T cells towards desired antigens of interest (*3*). We and others have used CAR technology to redirect Treg specificity to HLA-A2, finding that CAR expression significantly improves human Treg potency in humanised mouse models of transplantation (*4–9*). However, in fully immunocompetent mouse transplant models, HLA-A2-specific CAR (A2-CAR) Tregs do not induce indefinite graft survival (*10, 11*), suggesting that additional engineering may be required to fully unlock the protective capacity of these cells.

Programmed cell Death protein 1 (PD1) is an immune checkpoint molecule that is upregulated in T cells and Tregs following activation (*12*). In cancer research, multiple clinical trials have explored PD1 deletion as a strategy to limit T cell exhaustion (*13–16*), finding that PD1-deficient T cells are well-tolerated and may have an extended persistence compared to PD1-sufficient counterparts. In Tregs, evidence suggests that PD1 has a similarly negative role. Upon engagement of its ligands (PD-L1 or PD-L2), PD1 triggers intracellular signalling cascades that inhibit Treg activation (*7, 17, 18*) and consequently, subdue Treg function. High PD1 expression in human Tregs is correlated with a reduced suppressive ability and increased proinflammatory cytokine secretion in the contexts of cancer (*17*) and multiple sclerosis (*19*), and *in vitro* studies have shown that antibody-mediated blockade of PD1:PD-L1 interactions improves Treg function (*20–22*). Animal-based studies have also demonstrated that PD1:PD-L1 blockade promotes Treg proliferation/activity, thereby enhancing their ability to alleviate lupus-like disease (*23*), and selective removal of PD1 in FoxP3^+^ Tregs promotes the amelioration of experimental autoimmune encephalomyelitis and protects NOD mice from type 1 diabetes (*24*). Together, these observations suggest that removal of PD1 in therapeutic CAR Tregs may be beneficial for improved function.

Tregs suppress immune responses via multiple mechanisms, one of which is production of anti-inflammatory cytokines such as TGF-β and IL-10. However, amounts of IL-10 secreted by Tregs are relatively low (*25–27*), particularly in comparison to type 1 regulatory T (Tr1) cells that are defined by high IL-10 secretion (*28*). IL-10 primarily acts on myeloid cells to suppress proinflammatory cytokine production and limit the function of antigen presenting cells (APC), in turn suppressing the activation of adaptive immune cells (*29*). However, because of their low IL-10 production, both mouse and human Tregs are less able to suppress innate immune responses in comparison to Tr1 cells (*26, 27*), possibly resulting in suboptimal linked and/or bystander suppression which is thought to be critical for tolerance induction (*30–32*). Endowing Tregs with the ability to secrete high levels of IL-10 in an activation-dependent manner may overcome this limitation and enhance their therapeutic efficacy.

In this study, we explored whether the activation potential and immunosuppressive function of A2-CAR Tregs could be enhanced by ablating PD1 using CRISPR. In addition, we examined whether the endogenous *PDCD1* promoter could be exploited to control expression of an exogenous IL-10 transgene, thereby further improving the immunoregulatory function of A2-CAR Tregs.

## RESULTS

### Manufacturing fourth generation human CAR Tregs

In order to generate PD1-deficient A2-CAR Tregs, two gene editing procedures were performed: i) lentiviral transduction to deliver an HLA-A2-specific CAR (A2-CAR; Figure 1A), and ii) CRISPR editing to ablate PD1 (Figure 1B). Naïve CD4^+^CD25^hi^CD127^lo^CD45RA^+^CD45RO^−^ cells were sorted from the peripheral blood of healthy subjects and stimulated with artificial antigen presenting cells (aAPC) loaded with anti-CD3 (Figure 1C). After 24 hours, Tregs were transduced with a lentiviral vector that contained the open reading frames (ORF) of a previously characterised (*4, 7*) second-generation A2-CAR driven by an EF1α promoter, and a truncated nerve growth factor receptor (tNGFR) reporter driven by a minimal CMV promoter.

**Figure 1.**
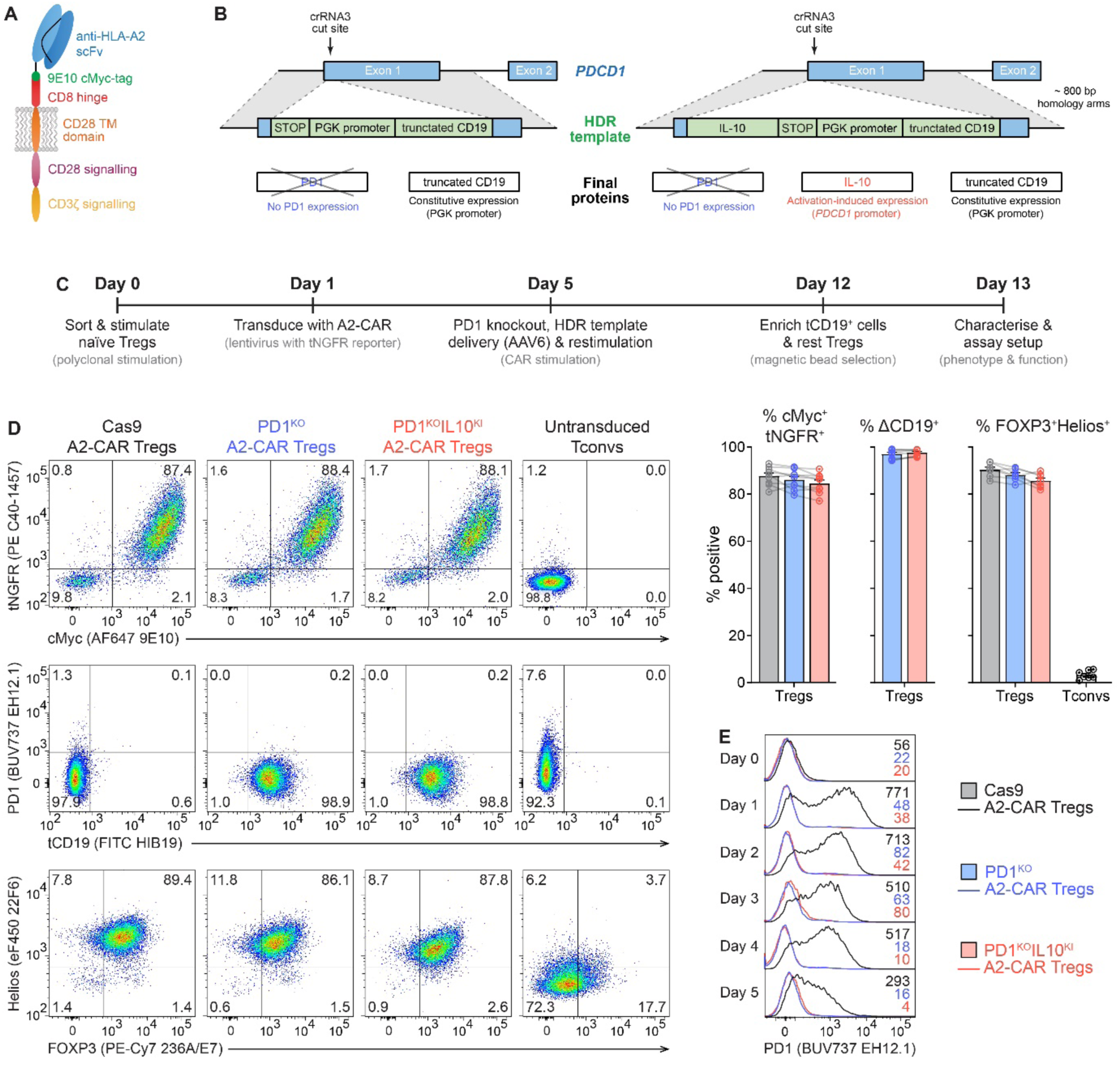
Generating PD1^KO^ and PD1^KO^IL10^KI^ A2-CAR Tregs. (**A**) Schematic diagram of the second-generation HLA-A2-specific CAR (A2-CAR). (**B**) Schematic diagram of the *PDCD1* locus and HDR templates. (**C**) Timeline for generating gene-edited A2-CAR Tregs. (**D**) Representative (left) and pooled (right) phenotypic characterisation of Tregs on day 13, gated on live CD4^+^ cells. Representative plots include % cells gated. (**E**) PD1 expression in A2-CAR Tregs following stimulation with HLA-A2^+^PD-L1^−^ K562 cells, with MFI values included. Gated on live CD4^+^cMyc^+^tNGFR^+^ and tCD19^+^ cells, where appropriate. Averaged data are mean + SEM with connected series representing individual subjects (n=3-9).

On day 5 of culture, Tregs were electroporated with Cas9 protein alone (control cells) or ribonuclear protein (RNP) complexes containing a *PDCD1*-specific gRNA that was confirmed to efficiently ablate PD1 (crRNA3; Supplemental Figure 1). For the latter conditions, a homology directed repair template (HDRT) was concurrently delivered by AAV6 transduction such that PD1-deficient A2-CAR Tregs could be identified by constitutive expression of a truncated CD19 (tCD19) reporter driven by a PGK promoter (Figure 1B, left). Alternatively, an HDRT containing a promoterless *IL10* ORF, in addition to the PGK-driven tCD19 reporter, was delivered into the *PDCD1* locus such that the endogenous cellular machinery that typically controls *PDCD1* would instead control expression of *IL10* (Figure 1B, right). Following electroporation, Tregs were restimulated with HLA-A2^+^ aAPCs to preferentially expand A2-CAR^+^ cells for an additional 7 days. On day 12 of culture, successfully edited tCD19^+^ cells were enriched using magnetic bead selection and rested overnight with a low-dose of IL-2 in preparation for assays that were performed on day 13 (Figure 1C).

Following expansion, >85% of the Tregs expressed the A2-CAR on their cell surface, as determined by staining for the cMyc tag present in the extracellular domain of the CAR construct (Figure 1D). These Tregs also maintained their expected constitutive expression of FOXP3 and Helios (Figure 1D). PD1-ablated Tregs were edited with a HDR efficiency of approximately 40% (tCD19: 45.2±15.0%, IL10tCD19: 40.2±14.4%, n=31) and enriched to a purity of >95% (Figure 1D). To confirm that the CD19^+^ A2-CAR Tregs were PD1-deficient, CAR-induced expression of PD1 was measured by flow cytometry. As expected, control Cas9 A2-CAR Tregs upregulated PD1, whereas no PD1 expression was observed for PD1^KO^tCD19^KI^ (herein referred to as PD1^KO^) or PD1^KO^IL10tCD19^KI^ (herein referred to as PD1^KO^IL10^KI^) A2-CAR Tregs (Figure 1E). Overall, these results demonstrate that human Tregs can be lentivirally transduced and CRISPR-edited without influencing their characteristic expression of FOXP3 and Helios.

### PD1-deletion increases A2-CAR Treg activation without compromising lineage stability

PD1 is a well-established negative regulator of T cell activation (*33*). To determine whether removal of PD1 in A2-CAR Tregs affects their state of activation following stimulation, Cas9 and PD1^KO^ A2-CAR Tregs were stimulated with HLA-A2^+^ aAPCs that co-expressed PD-L1, the ubiquitous PD1 ligand, and expression of Treg activation markers was measured after 48-hours. LAP and GARP (related to TGFβ secretion) were measured together with CTLA-4, an immunoregulation-related activation marker, and 4-1BB as a general indicator of cell activation. Compared to Cas9 A2-CAR Tregs, PD1^KO^ A2-CAR Tregs expressed significantly higher levels of all activation markers analysed, suggesting greater antigen sensitivity (Figure 2A).

**Figure 2.**
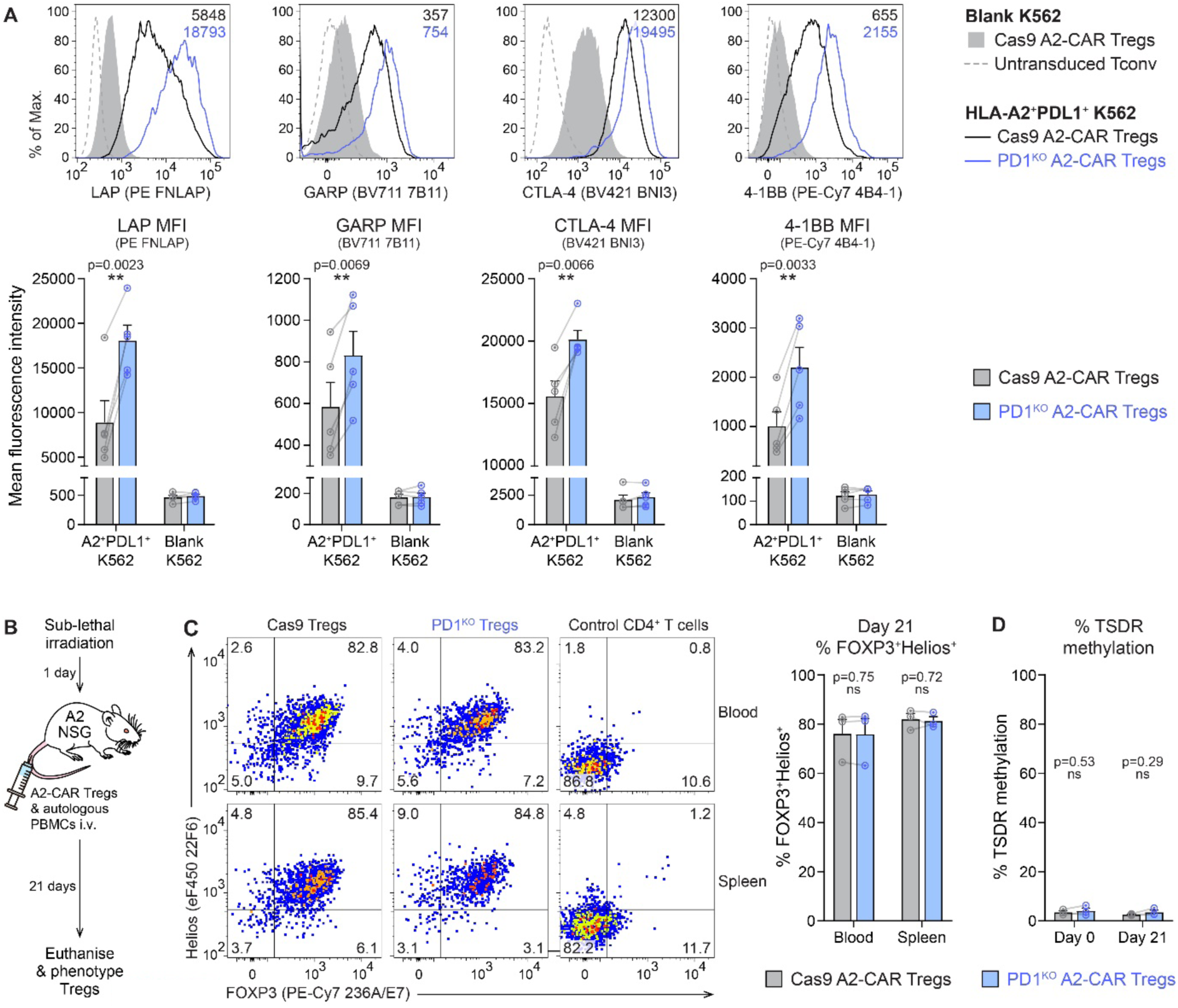
PD1-deficient Tregs express higher levels of activation markers and maintain their stability following CAR stimulation. (**A**) Expression of activation markers on Tregs following 48-hour co-culture with HLA-A2^+^PD-L1^+^ K562 cells. Gated on live CD4^+^cMyc^+^tNGFR^+^ cells and tCD19^+^ cells, where appropriate. (**B-D**) A2-CAR Tregs were intravenously co-administered with autologous PBMCs into transgenic A2-NSG mice. Mice were euthanized after 21-days. (**B**) Schematic diagram of model. (**C**) Expression of FOXP3 and Helios in hCD45^+^mCD45^−^hCD4^+^cMyc^+^ Tregs following euthanasia. Gates were set on hCD4^+^ cells that engrafted into mice reconstituted with PBMCs alone. (**D**) TSDR methylation of hCD45^+^mCD45^−^hCD4^+^cMyc^+^ Tregs isolated from splenocytes by cell sorting. Averaged data are mean + SEM with each connected series representing an individual subject (n=3-5). Statistical significance was determined using paired two-tailed Student’s t-tests with p-values shown.

It has previously been suggested that overstimulation of Tregs can drive instability and identity loss (*34*). To test whether PD1-deficient A2-CAR Tregs maintained their expression of FOXP3 and Helios following chronic CAR stimulation, cells were injected into NSG mice that ubiquitously expressed HLA-A2 (A2-NSG) for 3-weeks after which they were reisolated and analysed by flow cytometry (Figure 2B). Both Cas9 and PD1^KO^ A2-CAR Tregs maintained their expression of FOXP3 and Helios, indicating no loss of stability upon chronic stimulation (Figure 2C). This conclusion was further confirmed by analysing the methylation status of the *FOXP3* locus in human A2-CAR Tregs that were sorted from mouse spleens, which revealed no difference in methylation between Cas9 and PD1^KO^ cells (Figure 2D). Overall, these results demonstrate that PD1-ablation allows A2-CAR Tregs to achieve a higher level of activation upon CAR stimulation without impacting Treg lineage commitment.

### PD1^KO^IL10^KI^ A2-CAR Tregs display altered IL-10 secretion kinetics following CAR stimulation

In non-exhausted Tregs and T cells, PD1 is expressed in an activation-dependent manner (*35, 36*). To investigate whether the *PDCD1* promoter could be exploited to mediate activation-dependent expression of an exogenous transgene, an HDRT was designed to insert a promoterless *IL10* coding sequence into the *PDCD1* locus (Figure 1B, right). Due to the cut site of crRNA3, IL-10 protein synthesised from the *PDCD1* locus incorporated an MQIPQ N-terminal addition that was confirmed to not impact the immunosuppressive function of this cytokine (Supplemental Figure 2).

To assess whether PD1^KO^IL10^KI^ A2-CAR Tregs secreted IL-10 in response to CAR stimulation, A2-CAR Tregs were co-cultured with HLA-A2^+^PD-L1^−^ aAPCs cells for 72-hours after which cell culture supernatants were analysed for cytokine secretion. Treg cytokine secretion was compared to Tr1 cells, a population of induced immunosuppressive T cells characterised by high IL-10 secretion (*28*). PD1^KO^IL10^KI^ A2-CAR Tregs secreted ∼20-fold more IL-10 in response to CAR stimulation (p=0.0072), resulting in an average of 12.5 ng/mL which significantly exceeded that which was produced by Cas9 A2-CAR Tregs (p=0.0056) and A2-CAR Tr1 cells (p=0.029) seeded at the same cell concentration (Figure 3A). Further analysis of the relative quantities of cytokines produced revealed that the primary cytokine secreted by PD1^KO^IL10^KI^ A2-CAR Tregs was IL-10 (62.7% of total cytokine quantities analysed; Figure 3B). Conversely, whilst A2-CAR Tr1 cells also secreted high quantities of IL-10 (Figure 3A), they also produced significant levels of proinflammatory cytokines (e.g. IFNγ, IL-17A and TNFα), making the relative proportion of IL-10 secreted in comparison to proinflammatory cytokines low (only 1.9% of total cytokine quantities analysed; Figure 3B).

**Figure 3.**
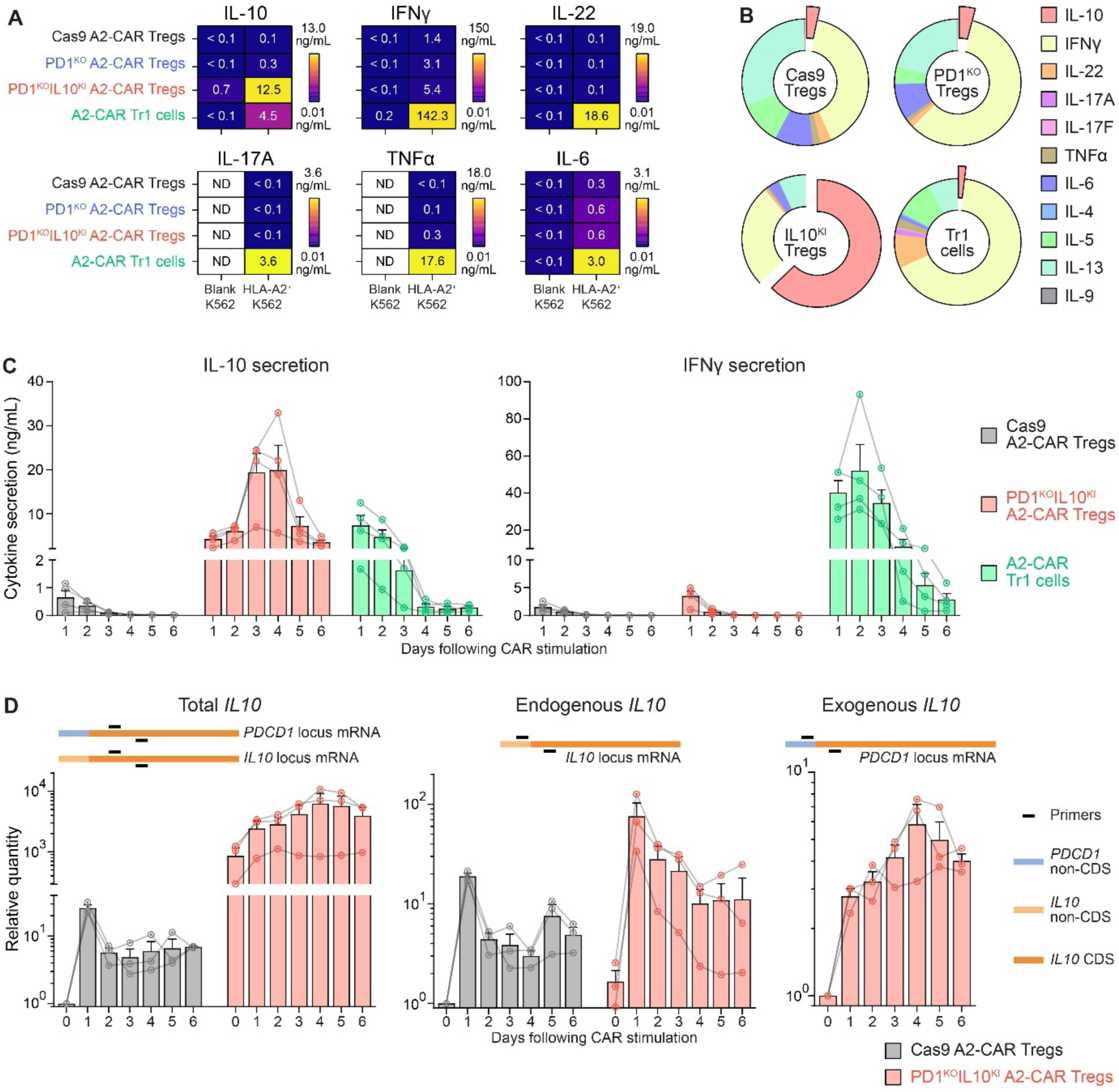
PD1^KO^IL10^KI^ A2-CAR Tregs secrete IL-10 with altered kinetics following CAR stimulation. (**A**) A2-CAR Treg and Tr1 cytokine secretion following 72-hour co-culture with blank or HLA-A2^+^PD-L1^−^ K562 cells. (**B**) Doughnut plot showing the amount of IL-10 secreted (red) as a proportion of 11 cytokines analysed. (**C**) A2-CAR Treg and Tr1 cytokine secretion profile following co-culture with HLA-A2^+^PD-L1^−^ K562 cells. Every 24-hours, culture supernatants were harvested, the cells were washed twice with PBS and re-cultured in fresh media with IL-2. (**D**) qPCR analysis of A2-CAR Treg *IL10* expression following co-culture with HLA-A2^+^PD-L1^−^ K562 cells. qPCRs were performed using primers that amplified mRNA corresponding to total *IL10* (left), endogenous *IL10* (middle) and exogenous *IL10* (right). Schematic diagrams showing primer recognition sites are provided. CDS = coding sequence. Averaged data are mean + SEM with each connected series representing an individual subject (n=3-5).

In addition to increasing the total amount of IL-10 secreted, control of *IL10* by the *PDCD1* promoter also has the potential to alter the kinetics of IL-10 secretion. To assess this, A2-CAR Tregs were stimulated with HLA-A2^+^ PD-L1^−^ aAPCs cells and every 24-hours, cell culture supernatants were harvested and Tregs were re-cultured in fresh media. This allowed cytokine secretion to be measured in 24-hour windows post-stimulation (Figure 3C and Supplemental figure 3). Control Cas9 A2-CAR Tregs secreted their highest quantities of IL-10 within the first 24-hours, and this steadily diminished over time. The quantities of IL-10 secreted by A2-CAR Tr1 cells were higher but with similar secretion kinetics to the Cas9 A2-CAR Tregs. In contrast, PD1^KO^IL10^KI^ A2-CAR Tregs secreted IL-10 with a unique kinetic profile that peaked approximately 72-hours post-stimulation. Importantly, the amount of IFNγ secreted by these cells remained low in comparison to Tr1 cells (Figure 3C and Supplemental Figure 3).

To further investigate the kinetics of *IL10* expression, qPCR analyses were performed to determine the amounts of mRNA synthesised from the *IL10* and *PDCD1* loci (Figure 3D). The kinetics of total *IL10* mRNA synthesis were comparable to the IL-10 secretion profiles seen in Figure 3C, with PD1^KO^IL10^KI^ A2-CAR Tregs producing more *IL10* than Cas9 A2-CAR Tregs that peaked 3-4 days post-stimulation. The majority of *IL10* expression was exogenous but interestingly, the endogenous *IL10* locus in the PD1^KO^IL10^KI^ A2-CAR Tregs was also more active, suggesting a positive feedback loop for IL-10 detection/synthesis, as has previously been described (*37–39*).

Overall, these results demonstrate that upon CAR stimulation, PD1^KO^IL10^KI^ A2-CAR Tregs secrete significantly greater quantities of IL-10 for a longer duration than control Tregs and Tr1 cells.

### PD1^KO^IL10^KI^ A2-CAR Tregs promote tolerogenic dendritic cells

IL-10 primarily acts on innate immune cells and has a key role in suppressing the antigen presenting function of dendritic cells (DC) (*29, 40–42*). To assess the effect of PD1^KO^IL10^KI^ A2-CAR Tregs on DC differentiation, CD14^+^ monocytes from HLA-A2^+^ individuals were co-cultured with A2-CAR Tregs in the presence of GM-CSF and IL-4 for 7-days after which the phenotype of the induced DCs was measured by flow cytometry (Figure 4A). PD1^KO^IL10^KI^ A2-CAR Tregs significantly promoted the expression of multiple markers associated with tolerogenic DCs including CD141, CD163, HLA-G and CD14 (Figure 4B). Interestingly, these DCs also upregulated CD80 but downregulated CD86, overall resulting in a phenotype that strongly resembled DC10 cells, an immunoregulatory population of DCs that are known to promote Tr1 differentiation and infectious tolerance (*38, 39*). In contrast, untransduced, Cas9 A2-CAR and PD1^KO^ A2-CAR Tregs all had a minor effect on influencing the DC differentiation process.

**Figure 4.**
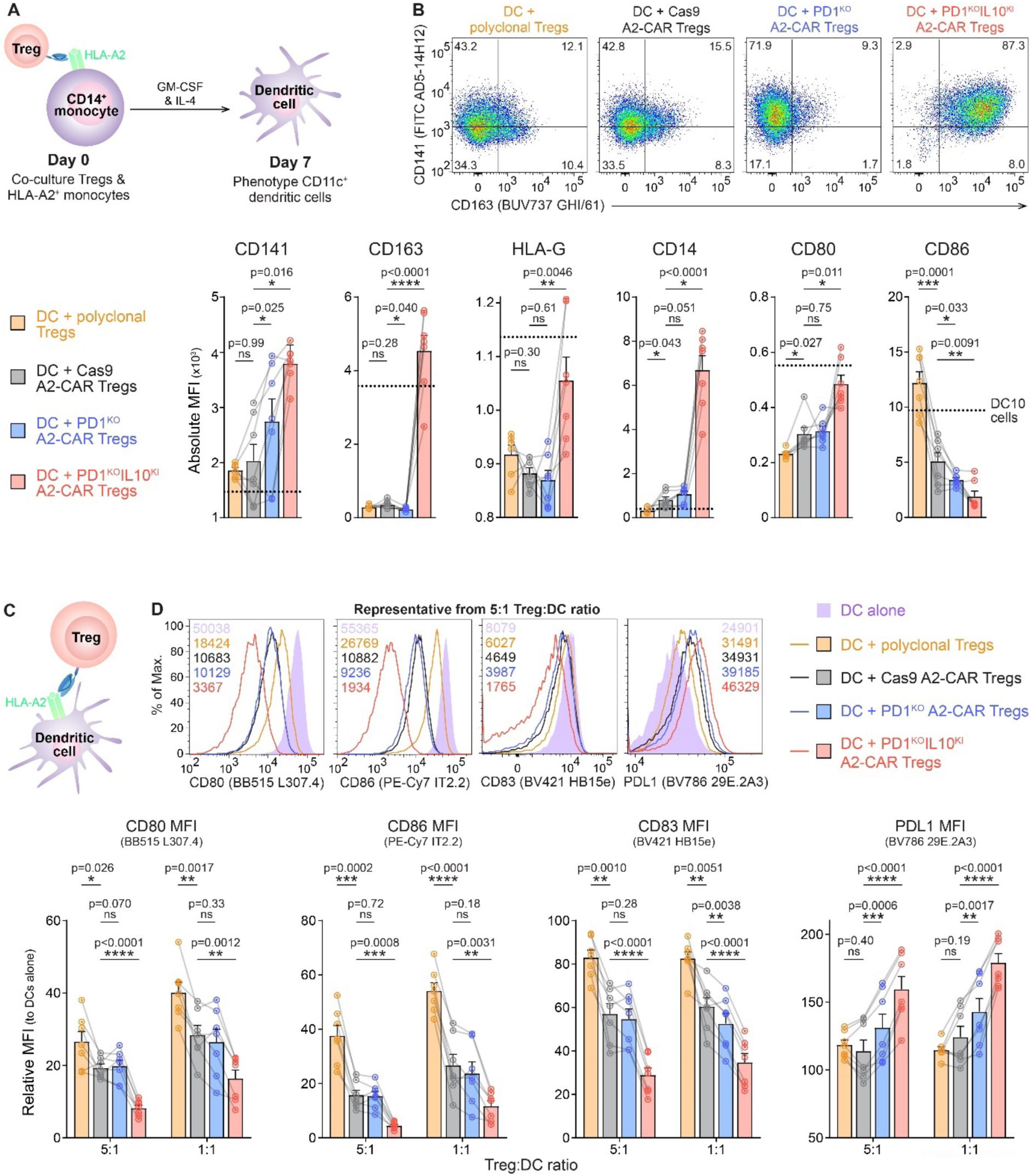
PD1^KO^IL10^KI^ A2-CAR Tregs induce a tolerogenic phenotype in dendritic cells. (**A-B**) CD14^+^HLA-A2^+^ cells were differentiated into dendritic cells with 100 ng/mL GM-CSF and 10 ng/mL IL-4, in the presence of Tregs (10:1 monocyte:Treg ratio). (**A**) Schematic diagram of experiment. (**B**) Expression of tolerogenic markers in live CD11c^+^CD4^−^ DCs was measured after 7-days. Representative plots include % cell gated (top) and pooled data shows absolute MFI values of indicated marker with dotted line representing DC10 cells (bottom). (**C-D**) Mature DCs were co-cultured with Tregs in the presence of 100 IU/mL IL-2 for 72-hours after which expression of co-stimulatory/inhibitory molecules was measured in live CD11c^+^CD4^−^ cells. (**C**) Schematic diagram of experiment. (**D**) DC phenotype after co-culture. Representative plots include MFI values (top) and pooled data shows relative MFI values compared to mDCs cultured alone (bottom). Averaged data are mean + SEM with each connected series representing an individual subject (n=7). Statistical significance was determined using paired two-tailed Student’s t-tests with p-values shown.

We next tested whether PD1^KO^IL10^KI^ A2-CAR Tregs could influence the antigen presenting capacity of established mature (m)DCs. Tregs were co-cultured with HLA-A2^+^ mDCs for 72-hours after which the DC phenotype was measured by flow cytometry (Figure 4C). A2-CAR expression significantly improved the ability of Tregs to suppress co-stimulatory molecules on the DCs, confirming our previous findings (*7*) (Figure 4D). Removal of PD1 in A2-CAR Tregs had a negligible effect on enhancing the suppression of co-stimulatory molecule expression but interestingly, this modification enabled the Tregs to induce PD-L1 expression on mDC, suggesting an enhanced capacity to promote tolerance. In contrast, PD1^KO^IL10^KI^ A2-CAR Tregs were significantly more suppressive than control Tregs, substantially inhibiting the expression of co-stimulatory molecules CD80, CD86 and CD83, as well as increasing expression of PD-L1 on mDCs. Overall, these results demonstrate that PD1^KO^IL10^KI^ A2-CAR Tregs acquire the ability to promote tolerogenic DC differentiation and have an enhanced capacity to suppress co-stimulatory molecule expression on proinflammatory mDCs.

### PD1^KO^IL10^KI^ A2-CAR Tregs have enhanced suppression of allo- and auto-reactive T cells

Tregs dampen deleterious immune responses by suppressing a range of cells including innate and adaptive immune cells. To investigate whether replacing PD1 with IL-10 improved the ability of Tregs to inhibit T cell proliferation, a variety of suppression assays were performed. Initial assays were performed in a polyclonal manner whereby Tregs were co-cultured with cell proliferation dye (CPD)-labelled responder PBMCs in the presence of anti-CD3/CD28 beads, such that Tregs were stimulated via the TCR rather than the CAR (Figure 5A). In this setup, Cas9 and PD1^KO^ Tregs inhibited CD4^+^ and CD8^+^ Tresponder proliferation to a similar degree whilst PD1^KO^IL10^KI^ Tregs were marginally more suppressive (Figure 5B). Further analysis of the CD19^+^ responder B cells revealed that PD1^KO^IL10^KI^ Tregs inhibited CD86 expression more than Cas9 or PD1^KO^ Tregs (Figure 5C), confirming that additional IL-10 expression enhances APC suppression, but also suggesting that the potential effects on T cells were not captured in this antigen-independent system (*40, 41*).

**Figure 5.**
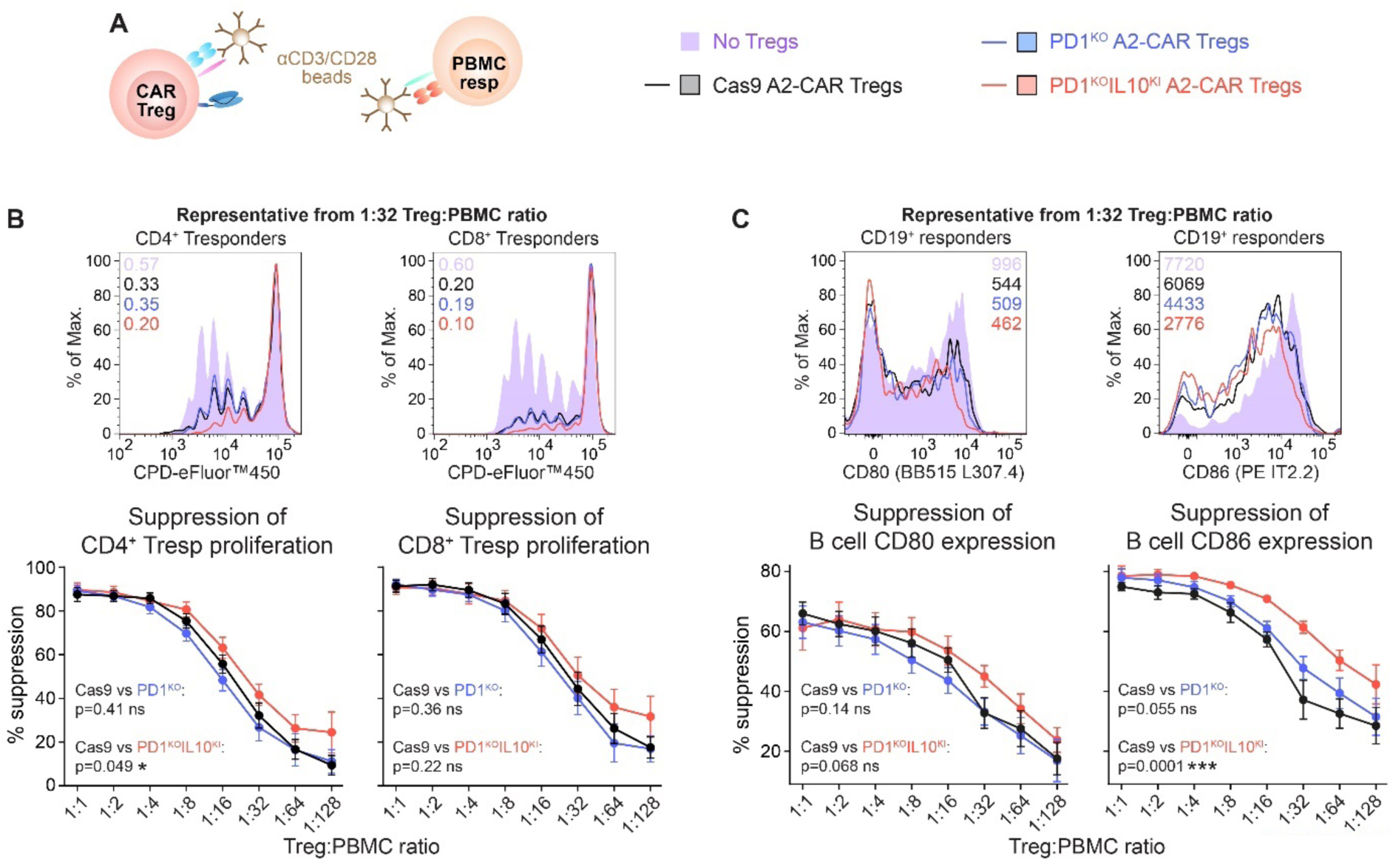
PD1^KO^IL10^KI^ Tregs have a minor functional advantage in an artificial polyclonal suppression assay setting. Tregs were co-cultured with CPD-eFluor™450-labelled PBMCs and anti-CD3/CD28 beads for 96-hours. (**A**) Schematic diagram of experiment. (**B**) CD4^+^ and CD8^+^ Tresponder proliferation (top; division indices provided) and % suppression of Tresponder proliferation, relative to PBMCs cultured alone (bottom). Gated on live CPD-eFluor™670^−^CPD-eFluor™450^+^CD4^+^ cells. (**C**) CD80 and CD86 expression in responder CD19^+^ B cells. Representative plots (top; MFI values provided) and % suppression of CD80 and CD86 relative to PBMCs cultured without Tregs (bottom), gated on live CD19^+^CD4^−^ cells. Averaged data are mean ± SEM (n=8). Statistical significance was determined using mixed-effects analyses with p-values shown.

We next tested the suppressive potency of PD1^KO^IL10^KI^ A2-CAR Tregs using two antigen- and APC-dependent suppression assays. In the first system, Tregs were co-cultured with HLA-A2^+^ DCs and the subsequent capacity of these DCs to stimulate direct alloreactive Tresponder cells was assessed (Figure 6A). A2-CAR expression significantly enhanced the ability of Tregs to inhibit alloreactive T cell proliferation (Figure 6B-C). However, PD1 deletion in the A2-CAR Tregs had a negligible effect, consistent with the finding that Cas9 and PD1^KO^ A2-CAR Tregs equally suppressed co-stimulatory molecule expression in mDCs (Figure 4D). In contrast, PD1^KO^IL10^KI^ A2-CAR Tregs were significantly more effective at suppressing alloreactive CD4^+^ T cell proliferation than all other Treg conditions analysed (Figure 6B-C).

**Figure 6.**
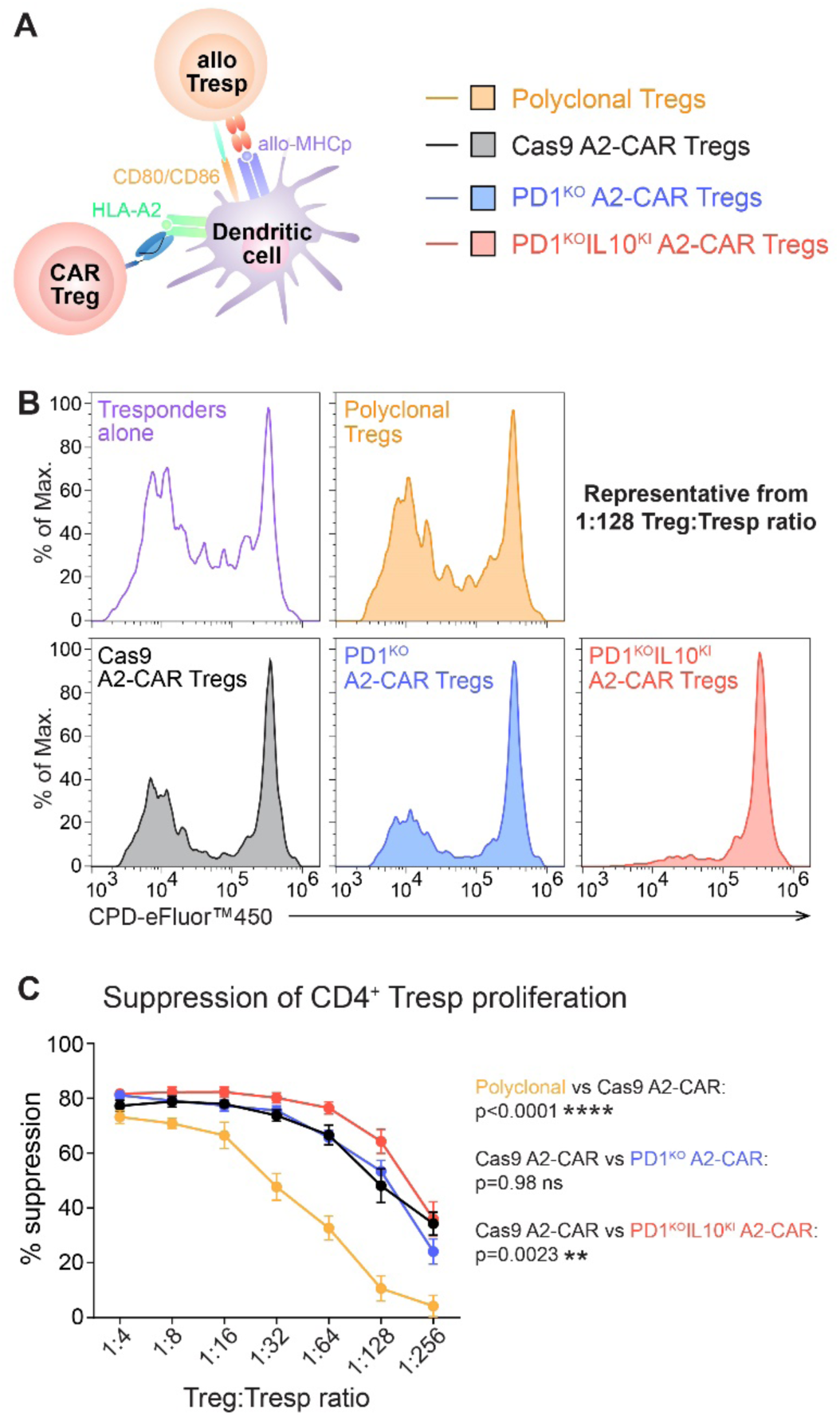
PD1^KO^IL10^KI^ A2-CAR Tregs suppress alloantigen-specific T cell proliferation. HLA-A2^+^ DCs were co-cultured with HLA-A3^−^ Tregs in the presence of 100 IU/mL IL-2 for 72-hours. Allogeneic CPD-eFluor™450-labelled HLA-A3^+^CD3^+^ responder T cells were then added and co-cultures were maintained for an additional 96-hours. (**A**) Schematic diagram of experiment. (**B**) Representative CD4^+^ Tresponder proliferation, gated on live HLA-A3^+^CPD-eFluor™670^−^CPD-eFluor™450^+^CD4^+^ cells. (**C**) % suppression of Tresponder proliferation, relative to Tresponders stimulated without Tregs. Averaged data are mean ± SEM (n=10). Statistical significance was determined using mixed-effects analysis with p-values shown.

To further confirm the enhanced suppressive function of PD1^KO^IL10^KI^ A2-CAR Tregs, an alternative APC-dependent suppression assay was performed in which the ability of these cells to inhibit autoreactive islet-specific CD4^+^ T cells was measured. CD4^+^ Tresponder cells were transduced to express an islet autoantigen-specific TCR, specifically the 4.13-TCR (*30*), which recognises glutamic acid decarboxylase (GAD)65 peptide presented in the context of HLA-DR4 (Figure 7A).

**Figure 7.**
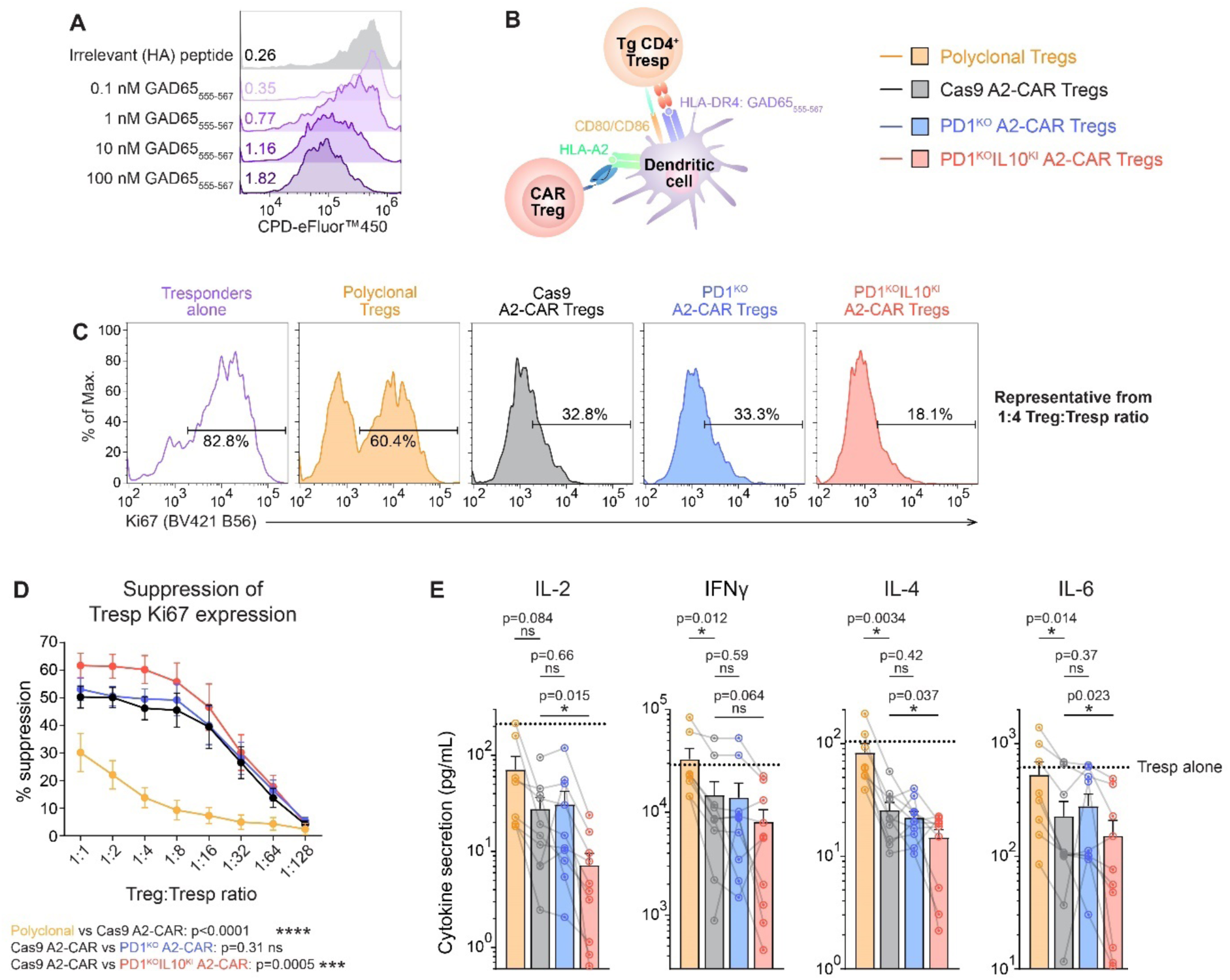
PD1^KO^IL10^KI^ A2-CAR Tregs suppress islet-specific T cells. HLA-A2^+^HLA-DR4^+^ DCs were co-cultured with Tregs for 48-hours in the presence of 100 IU/mL IL-2. CD4^+^4.13-TCR^+^ Tresponders and 1 nM GAD65 peptide were then added and co-cultures were maintained for an additional 48-hours. (**A**) Proliferation of CD4^+^4.13-TCR^+^ T cells following 96-hour co-culture with immature HLA-DR4^+^ DCs and varying concentrations of GAD65 peptide (without Tregs). Control conditions were pulsed with 100 nM of an irrelevant HA peptide. Division indices are provided. (**B**) Schematic diagram of suppression assay setup. (**C**) Representative CD4^+^ Tresponder proliferation, gated on live CD4^+^CPD-eFluor™670^−^mTCRβ^+^ cells. (**D**) % suppression of Tresponder proliferation, relative to Tresponders stimulated without Tregs. (**E**) Cytokine analysis from 1:128 Treg:Tresp ratio. Dotted lines represent cultures without Tregs. Averaged data are mean ± SEM (n=10). Statistical significance was determined using mixed-effects analysis (**D**) or paired two-tailed Student’s t-tests (**E**) with p-values shown.

A2-CAR Tregs were co-cultured with HLA-A2^+^HLA-DR4^+^ DCs and the subsequent ability of these DCs to stimulate 4.13-TCR^+^ Tresponder cells in the presence of GAD65 peptide was assessed (Figure 7B). CD4^+^ Tresponder proliferation was measured by the induced expression of Ki67, a well-established proliferation marker. Similar to observations with alloreactive T cells, Cas9 and PD1^KO^ A2-CAR Tregs exhibited a comparable level of suppression whilst PD1^KO^IL10^KI^ A2-CAR Tregs inhibited Ki67 expression in autoreactive Tresponder cells significantly more effectively (Figure 7C-D). Analysis of the supernatants from these suppression assays also revealed that cultures containing PD1^KO^IL10^KI^ A2-CAR Tregs had significantly less pro-inflammatory cytokines (Figure 7E). Overall, these results show that PD1^KO^IL10^KI^ A2-CAR Tregs are superior to A2-CAR Tregs (with or without PD1) and have an enhanced ability to suppress allo- and auto-reactive T cells.

## DISCUSSION

In this study, we built on the concept of using CAR Tregs as an immunoregulatory cell therapy product and created so-called “fourth generation” CAR Tregs that were engineered to simultaneously remove an inhibitory signal and enhance a suppressive mechanism. Using PD1 as a representative inhibitory molecule and IL-10 as a prototypical suppressive cytokine, we demonstrate that the function of CAR Tregs can be substantially increased by replacing *PDCD1* with *IL10*. Working in the human system with an A2-CAR, we found that PD1^KO^IL10^KI^ A2-CAR Tregs took on advantageous properties of Tr1 cells, adopting an enhanced ability to drive tolerogenic DC development and control both allo- and auto-reactive T cells. Importantly, this was achieved without the deleterious secretion of pro-inflammatory cytokines, a potential limitation of Tr1 cells (*27, 43*). These findings are the first to demonstrate the benefits of using genome engineering to enhance a functional pathway in CAR Tregs and create cells with “hybrid” functions that are usually present in two distinct cell types.

Seeking antigen-dependent control of a functionally advantageous transgene, we chose to target the *PDCD1* locus as PD1 expression is activation induced (*12*) and there is the added benefit that PD1 signalling is deleterious for Treg activation (*7, 17, 18*). Although some studies found that PD1 signalling promotes FOXP3 expression and conversion of T cells into Tregs (*44–47*), most studies have determined that PD1 signalling in established Tregs is detrimental for their function (*7, 17, 19, 23, 24*). In our study, CRISPR-mediated PD1 ablation increased the activation potential of human A2-CAR Tregs but, unlike cancer-relevant CAR-T cells (*48, 49*), this modification did not significantly improve their function. Importantly however, PD1 ablation did not compromise Treg stability or diminish their function in any DC or T cell directed assay.

Multiple Treg mechanisms of action have been described, but the relevance of each pathway in different disease contexts remains relatively uncharacterised. We selected IL-10 as an ideal candidate for enhancing the suppressive function of Tregs as it is a well-established immunoregulatory cytokine that is secreted at relatively low levels compared to Tr1 cells (*27*). IL-10 therapy has been investigated clinically as a monotherapy (*50, 51*) but its short half-life is thought to have impacted the levels of IL-10 present in the required tissues (*52*), limiting its efficacy and suggesting that *in situ* secretion at sites of inflammation may be more advantageous. IL-10 primarily achieves its immunosuppressive effect by acting on myeloid cells; it can act directly on T cells but the consequences of this are less significant since IL-10 signalling counters co-stimulatory signals, but not TCR or IL-2-mediated signalling cascades (*40, 41*). Furthermore, unlike myeloid cells, naïve and resting T cells express low levels of the IL-10 receptor α-chain which is responsible for exclusively binding IL-10 (*42*). These findings offer an explanation for the modest functional advantage the PD1^KO^IL10^KI^ A2-CAR Tregs displayed in an APC-deficient suppression assay system (Figure 5).

A significant advantage of editing CAR Tregs to express IL-10 instead of using CAR Tr1 cells is the low amounts of pro-inflammatory cytokines produced by the former. As a consequence, the concept of endowing CAR Tregs with the ability to constitutively express IL-10 has previously been explored, with mixed results (*53, 54*). In these studies, the CAR and *IL10* were delivered in the same viral-vector-based constructs. Constitutive IL-10 expression was found to increase the suppressive function of human A2-CAR Tregs (*53*) but did not have an effect on mouse CAR Tregs redirected towards factor VIII (*54*). Furthermore, Tregs constitutively secreting could be deleterious *in vivo* as they may promote pan-immunosuppression and possibly inflammation given the paradoxically pleiotropic effects of IL-10 on driving CD8^+^ T cell expansion and B cell antibody secretion (*55–58*). Our strategy reduces this concern as IL-10 is secreted in an activation-dependent manner, primarily at sites of inflammation where it is required. Emerging protein engineering strategies that have been used to improve the binding of IL-10 to its cognate receptor may also be adapted to reduce these unwanted inflammatory side effects of IL-10 (*59, 60*). Other immunoregulatory or tissue reparative molecules could similarly be incorporated into the *PDCD1* or other antigen-stimulated loci to improve CAR Treg function.

Given that PD1 is transiently expressed upon stimulation in non-exhausted cells, various cancer immunotherapy studies have explored the advantages of exploiting the *PDCD1* locus to regulate expression of exogenous transgenes. Insertion of an anti-CD19 CAR into the *PDCD1* locus of T cells using a non-viral gene editing strategy was shown to reduce CAR-T cell exhaustion upon repetitive *in vitro* stimulation (*61*) and the therapeutic efficacy of these cells was recently confirmed in a phase I clinical trial (*16*). In our study, we modified this strategy to insert a functionally-relevant transgene into the *PDCD1* locus, yielding a Treg product that constitutively expressed an HLA-A2-specific CAR and expressed the immunosuppressive cytokine IL-10 in response to antigen stimulation. Using a similar gene editing strategy, Kim *et al*. also observed activation-dependent expression in their cancer-based study that inserted IL-12 into the *PDCD1* locus of TCR-edited NY-ESO T cells (*36*).

The clinical implications of this study were impacted by the lack of a suitable humanised *in vivo* model in which we could test the enhanced suppressive capacity of PD1^KO^IL10^KI^ A2-CAR Tregs, compared to control Cas9 A2-CAR Tregs. We considered use of the xenogeneic graft-versus-host disease (xenoGVHD) model, where we previously showed the heightened suppressive function of A2-CAR Tregs compared to control CAR Tregs (*4, 7*), but in this model IL-10 is known to counterintuitively promote xenoGVHD (*62, 63*), likely due to proinflammatory effects of IL-10 on CD8^+^ T cells and the lack of APCs in these models (*64*). Moreover, the antigen presenting function of DCs is notoriously absent in PBMC-reconstituted NSG-mice (*65, 66*). Thus, to the best of our knowledge, there is currently no humanised mouse model in which the predominantly APC-mediated effects of PD1^KO^IL10^KI^ A2-CAR Tregs could be studied. Another consideration is the logistical complexity of introducing three genetic modifications using lentivirus and CRIPSR editing. We opted to use AAV6 to deliver an HDRT encoding the IL-10 transgene into Tregs, but future studies should consider using non-viral gene delivery systems to decrease the cost, complexity and potential immunogenicity of AAV6 (*61, 67, 68*).

Overall, this study shows the potential of gene engineering to enhance the therapeutic potency of CAR Tregs. Although IL-10 secretion is often cited as a key Treg mechanism of action, critically important for immune regulation in tissues at environmental interfaces (*69*), its expression is low in human Tregs isolated from blood (*27*). Thus, blood-derived Tregs tested in clinical trials to date have likely had low IL-10 expression. Our data show that this pathway can be easily introduced ectopically to significantly improve Treg function, without loss of lineage stability. The findings set the stage for using this strategy to make Tregs with enhanced therapeutic effects for multiple applications in transplantation and autoimmunity.

## MATERIALS & METHODS

### Study design

The objectives of this study were to address the following pre-determined hypotheses: i) CRISPR-mediated removal of PD1 would allow human CAR Tregs to achieve a higher activation state in response to a specific CAR stimulant, ii) the *PDCD1* locus could be exploited to facilitate activation-induced expression of an IL-10 transgene and iii) replacing *PDCD1* with *IL10* would enhance the suppressive potency of human CAR Tregs. Experiments were performed using cells isolated from the peripheral blood of anonymized human subjects. Male subjects were selected for Treg stability experiments in which TSDR analyses were performed (Figure 2). For all other experiments, subjects were not stratified according to sex. Within each experiment, Cas9, PD1^KO^ and PD1^KO^IL10^KI^ A2-CAR Tregs (and A2-CAR Tr1 cells) were generated from the same subject to reduce donor-to-donor variability; the relative contribution of each subject is shown with connected points. Experiments were performed at least twice with 2-4 individual subjects per experiment; the number of biological replicates is stated in the figure legends and sample sizes were established prior to performing the experiments, based on our previous experience with similar experimental approaches to achieve statistical significance. Cytokine secretion values were regarded as “not detectable” and excluded from analyses if the predicted concentrations were below the LEGENDPlex™ (BioLegend) threshold of detection or consistently < 1 pg/mL for multiple donors; no other data were excluded from our analyses.

### Subjects, mice & ethical approvals

Peripheral blood in the form of buffy coat products was obtained from anonymized healthy adults via Canadian Blood Services (Vancouver BC, Canada) with informed consent and ethical approval from The University of British Columbia Clinical and Canadian Blood Service Research Ethics Boards (H18-02553).

Transgenic NOD.Cg-*Prkdc^scid^ Il2rg^tm1Wjl^*/SzJ mice that ubiquitously express HLA-A2 (A2-NSG; Jackson Laboratory) were bred in-house and maintained under specific pathogen free conditions. All *in vivo* experiments were carried out in accordance with the National Research Council’s Guide for the Care and Use of Laboratory Animals using protocols approved by The University of British Columbia Animal Care Committee (A22-0120).

### Molecular biology & vector synthesis

Homology directed repair templates (HDRT) were designed with ∼800 bp homology arms to insert designated sequences into the *PDCD1* locus following CRISPR-mediated cutting with crRNA3. An *IL10-tCD19* ORF was gene synthesised (GeneArt) and cloned by Gibson assembly (NEB) into an AAV6 transfer plasmid (*70*), thereby generating pssAAV_PDCD1cr3.HDRT-IL10tCD19 (Figure 1B). The IL10-encoding sequence was then removed to generate pssAAV_PDCD1cr3.HDRT-tCD19.

pCCL_A2CAR-tNGFR and pCCL_HLAA2-eGFP were generated as previously described (*4*). pCCL_wtIL10-tNGFR and pCCL_MQIPQ-IL10-tNGFR were generated by amplifying *IL10* from the pssAAV_PDCD1cr3.HDRT-IL10tCD19 vector and using Gibson assembly to insert this PCR product into the pCCL_A2CAR-tNGFR vector. pELNS_PDL1 was generated by cloning PCR-amplified *PDL1* cDNA (Sino Biological) into the pELNS vector using Gibson assembly. pCCL_4.13TCR-tNGFR was generated by replacing the *A2CAR* ORF in the pCCL_A2CAR-tNGFR vector with 4.13TCRα and 4.13TCRβ-chain sequences (*30*) that were split by a P2A sequence. TCR constant regions were replaced with murine TRAC and TRBC sequences to allow identification of transduced cells.

All vectors were validated by restriction digest and Sanger sequencing before use.

### Cell line culture

HEK293T clone 17 cells (ATCC) were cultured in IMDM supplemented with 10% foetal bovine serum (FBS), 100 U/mL penicillin-streptomycin and 2 mM GlutaMAX™ (all from Thermo Fisher Scientific). L cells expressing human CD32, CD58 and CD80 (*71*) were cultured in RPMI (Thermo Fisher Scientific) supplemented with 10% foetal bovine serum (FBS; Hyclone Laboratories Inc.), 100 U/mL penicillin-streptomycin and 2 mM GlutaMAX™. HLA-A2^+^ L cell derivatives were generated by lentivirally transducing CD32^+^CD58^+^CD80^+^ L cells with pCCL_HLAA2-eGFP. Cells were passaged by washing with PBS, incubating with 0.05% Trypsin-EDTA (Thermo Fisher Scientific) for 2-3 minutes and neutralising with cell culture media.

K562 cells (ATCC) were cultured in RPMI supplemented with 10% FBS, 100 U/mL penicillin-streptomycin and 2 mM GlutaMAX™. HLA-A2^+^PD-L1^−^ and HLA-A2^+^PD-L1^+^ K562 derivatives were generated by lentivirally transducing blank K562s with pCCL_HLAA2-eGFP and pELNS_PDL1.

### Virus production

To generate lentiviral particles, HEK293T/17 cells were transfected with either i) pCCL_A2CAR-tNGFR, ii) pCCL_4.13TCR-tNGFR, iii) pCCL_HLAA2-eGFP, or iv) pELNS_PDL1 and a mixture of pRSV-REV, pMDLG/pRRE, pMD2.g and pAdVAntage™ Vector (further details on addgene.org) using calcium phosphate (made in-house). Cell supernatants were harvested 45-48 hours post-transfection and lentiviral particles were concentrated by ultracentrifugation at 76,755 g. Viral titres were calculated by limiting dilution transduction of HEK293T/17 cells and virus aliquots were stored at –80°C.

To generate AAV6 viral particles, HEK293T/17 cells were co-transfected with either pssAAV_PDCD1cr3.HDRT-tCD19 or pssAAV_PDCD1cr3.HDRT-IL10tCD19 and a mixture of pHelper and pAAV6-Rep-Cap (both from Cell Biolabs) using calcium phosphate. Culture supernatants were replenished after 12-16 hours and harvested ∼68-hours post-transfection. AAV6 particles were concentrated using the AAVpro Purification Kit, quantified using the AAVpro quantification kit (both from Takara Bio) and stored at –80°C.

### RNP generation

*PDCD1*-targeting crRNA1 (5’-CGTCTGGGCGGTGCTACAACTGG-3’) (*72–74*), crRNA2 (5’-GGCCAGGATGGTTCTTAGGTAGG-3’) (*73, 75*) and crRNA3 (5’-CGACTGGCCAGGGCGCCTGTGGG-3’) (*49, 76–78*) were reconstituted at 200 μM in nuclease-free Duplex buffer and duplexed at a 1:1 molar ratio with tracrRNA (all from IDT) by heating to 95°C for 5 minutes and cooling to room temperature, as previously described (*70*). Resulting gRNA was then combined at a 2:1 molar ratio Cas9-NLS protein (QB3 Macrolab) and incubated for 10 minutes at room temperature to generate RNPs. RNPs were used immediately or stored at –80°C. Each electroporation contained < 2×10^5^ cells and 20 pmol Cas9 ± 40 pmol gRNA in a total volume of 10 μL.

### Treg isolation, gene editing & expansion

CD4^+^ T cells were enriched from peripheral blood of HLA-A2^−^ subjects using the Rosettesep™ Human CD4^+^ T Cell Enrichment Cocktail (STEMCELL Technologies) and CD25^+^ cells were separated using CD25 MicroBeads II (Miltenyi Biotec). Naïve CD4^+^CD25^hi^CD127^lo^CD45RA^+^CD45RO^−^ Tregs and naïve CD4^+^CD25^−^CD127^+^CD45RA^+^CD45RO^−^Tconvs were then purified by cell sorting. Sorted T cells were stimulated with artificial APCs (L cells expressing human CD32, CD58 and CD80; irradiated with 7500 cGy) loaded with anti-CD3 mAb (OKT3, UBC AbLab) and cultured in ImmunoCult™ XF (STEMCELL Technologies) supplemented with 100 U/mL penicillin-streptomycin. Tregs and Tconvs were cultured with 1,000 IU/mL and 100 IU/mL recombinant human IL-2, respectively (Proleukin®, Prometheus Laboratories Inc.).

Tregs were transduced with the A2-CAR construct 24-hours post-stimulation using lentivirus at an MOI of 10, as previously described (*4*). On day 5, Tregs were CRISPR-edited using the Neon Transfection System 10 µL Kit (Thermo Fisher Scientific) and restimulated, as previously described (*70*). Briefly, Tregs were harvested, washed twice with PBS and resuspended in Buffer T (< 2×10^5^ cells per 10 μL) containing Cas9 or complexed RNPs. Cells were then electroporated in one pulse with 1400 V for 30 ms using the Neon NxT Electroporation System and immediately transferred into cell culture wells containing prewarmed antibiotic-free ImmunoCult™ XF, 1,000 IU/mL IL-2, AAV6 (35-45,000 vg/cell) and HLA-A2^+^ L cells (without OKT3). Tregs were maintained in culture for 7 days after electroporation.

On day 12 of culture, edited tCD19^+^ cells were enriched using the EasySep™ Human CD19 Positive Selection Kit II (STEMCELL Technologies). All cells were rested overnight by culturing in the presence of 100 IU/mL (Tregs) or 10 IU/mL (Tconvs) IL-2 and assays were performed on day 13 of culture.

### Tr1 cell isolation, culture and transduction

Tr1 cells were isolated using an IL-10 capture approach, as previously described (*27*). Briefly, enriched CD4^+^ T cells were stimulated with Dynabeads™ Human T-Expander CD3/CD28 (1:16 bead:cell ratio; Thermo Fisher Scientific) overnight, cells primed to secrete IL-10 were stained using the IL-10 Secretion Assay Detection Kit (Miltenyi Biotec) and IL-10^+^ cells were isolated by cell sorting. Isolated Tr1 cells were stimulated with OKT3-loaded L cells, cultured in the presence of 100 IU/mL IL-2, transduced to express the A2-CAR, re-stimulated with HLA-A2^+^ L cells and rested as described above, in parallel with Tregs from matched donors.

### Flow cytometry & cell sorting

Cells were stained using previously described protocols (*79, 80*) in PBS supplemented with 1% bovine serum and 5 mM EDTA (Sigma-Aldrich) with fluorescently-conjugated antibodies specific for mouse CD45 (PE-Cy7 30-F11; Thermo Fisher Scientific or BUV395 30-F11; BD Biosciences) and human CD45 (V500 HI30), CD4 (V500 RPA-T4, BV605 RPA-T4, APC-R700 SK3 or BV711 SK3), CD271 (PE C40-1457) GARP (BV711 7B11), CD8a (PE HIT8a), PD1 (BUV737 or PE-Cy7 EH12.1), CD19 (FITC HIB19), CD80 (BB515 L307.4), CD83 (APC HB15e), CD86 (PE IT2.2), CD163 (BUV737 GHI/61) (all from BD Bioscience), CD4 (FITC RPA-T4), CD45RA (PE-Cy7 HI100), CD45RO (eF450 UCHL1), FOXP3 (PE-Cy7 236A/E7), LAP (PE FNLAP), CD4 (FITC RPA-T4), CD8a (eF450 SK1), Helios (eFluor™450 22F6), CD86 (PE-Cy7 IT2.2), Ki67 (BV421 B56), CD11c (BUV805 3.9), HLA-G (PerCP-eF710 87G), HLA-A3 (FITC GAP.A3) (all from Thermo Fisher Scientific), CD4 (BV421 OKT4), PD1 (BV786 EH12.1), CTLA-4 (BV421 BNI3), 4-1BB (PE-Cy7 4B4-1), PD-L1 (BV785 29E.2A3), HLA-A2 (PE BB7.2) (all from BioLegend), CD25 (PE 4E3), CD141 (FITC AD5-14H12) (all from Miltenyi Biotec), CD127 (APC-AF700 R34.34; Beckman Coulter) and cMyc (AF647 9E10; UBC AbLab).

Dead cells were excluded using Fixable Viability Dye (FVD) eFluor™ 780. For samples containing APCs, Fc receptors were blocked by incubating with a polyclonal Fc block for 10 minutes at room temperature before surface staining. Staining for intracellular markers was performed using the Foxp3/Transcription Factor Staining Buffer Set. Cell proliferation was measured using Cell Proliferation Dye (CPD) eFluor™450 or 670 (all from Thermo Fisher Scientific).

Cells were sorted using a MoFlo® Astrios (Beckman Coulter) or FACSAria™ Fusion (BD Biosciences). Flow cytometry data were acquired using an LSRFortessa™ II, FACSymphony™ A5, FACSymphony™ A1 (all from BD Biosciences) or CytoFLEX (Beckman Coulter) and analysed using FlowJo X (BD Biosciences).

### Activation assays, IL-10 secretion analyses & *IL10* qPCR

Rested Tregs were co-cultured with blank, HLA-A2^+^PD-L1^−^ or HLA-A2^+^PD-L1^+^ K562 cells (5:1 Treg:K562 ratio) in ImmunoCult™ XF supplemented with 100 U/mL penicillin-streptomycin and 100 IU/mL IL-2. Tr1 cells were similarly treated in cultures supplemented with 10 IU/mL IL-2. Induced expression of activation markers was determined by flow cytometry after 48-hours. For cytokine secretion analyses, cell culture supernatants were harvested after 72-hours. Alternatively, cytokine secretion kinetics were determined by collecting supernatants every 24-hours; after each collection, cells were washed twice with PBS and re-cultured in fresh media supplemented with IL-2. Cytokines were measured by LEGENDPlex™ using a 12-plex Human Th Cytokine Panel (BioLegend).

For qPCR analyses, cells were harvested at 24-hour timepoints following stimulation, washed with PBS and lysed with Buffer RLT (Qiagen) supplemented with 1% β-mercaptoethanol (Sigma-Aldrich). RNA was extracted using the RNeasy Micro Kit (Qiagen) and reverse transcribed using qScript™ cDNA SuperMix (QuantaBio). qPCR analyses were performed using PerfeCTa® SYBR® Green FastMix®, Low ROX™ (QuantaBio) and run on a ViiA™ 7 Real-Time PCR System (Applied Biosystems, Thermo Fisher Scientific). Data were analysed using a 2^−ΔΔCt^ approach with 18S and β2-microglobulin as housekeeping controls and the following primers: total *IL10* (FWD 5’-GCTCAGCACTGCTCTGTTGCCTG-3’, REV 5’-CTCGAAGCATGTTAGGCAGGTTGCC-3’), endogenous *IL10* (FWD 5’-CAGACTTGCAAAAGAAGGCATGCAC-3’, REV 5’-CTCGAAGCATGTTAGGCAGGTTGCC-3’), exogenous *IL10* (FWD 5’-GTGGAGAAGGCGGCACTCTGGTG-3’, REV 5’-CTCGAAGCATGTTAGGCAGGTTGCC-3’), 18S (FWD 5’-CAAGACGGACCAGAGCGAAA-3’, REV 5’-GGCGGGTCATGGGAATAAC-3’) and β2-microglobulin (FWD 5’-ATGTCTCGCTCCGTGGCCTTAG-3’, REV 5’-CCATTCTCTGCTGGATGACGTGA-3’).

### CD14^+^ cell isolation & dendritic cell differentiation

CD14^+^ cells were isolated from PBMCs using the EasySep™ Human CD14 Positive Selection Kit II (STEMCELL Technologies) to a purity of > 97%, as confirmed by flow cytometry. Immature and mature DCs were differentiated as previously described (*7, 81*). Briefly, CD14-enriched cells were resuspended in ImmunoCult™ XF supplemented with 100 U/mL penicillin-streptomycin and cultured in the presence of 50 ng/mL GM-CSF and 100 ng/mL IL-4 (both from STEMCELL Technologies) for 7-days, yielding immature DCs. To mature DCs, cultures were additionally supplemented with 10 ng/mL IL-1β, 100 ng/mL IL-6 (both from STEMCELL Technologies), 50 ng/mL TNFα (eBioscience) and 1 μg/mL prostaglandin E2 (Tocris) from day 5, as well as 50 ng/mL IFNγ (eBioscience) from day 6.

For assays in which DCs were differentiated in the presence of Tregs, CD14^+^ cells from an HLA-A2^+^ individual were co-cultured with Tregs (10:1 monocyte:Treg ratio) in ImmunoCult™ XF supplemented with 100 U/mL penicillin-streptomycin and 5% human serum (Wisent Bioproducts) in the presence of 100 ng/mL GM-CSF and 10 ng/mL IL-4 for 7 days. Positive control cell cultures (DC10 cells) were additionally supplemented with 10 ng/mL IL-10 (STEMCELL Technologies). The phenotype of the differentiated DCs was then measured by flow cytometry, gating on live CD11c^+^ cells.

### IL-10 functional testing

The immunosuppressive function of IL-10 variants was tested as previously described (*79, 82*). Wild-type and MQIPQ-modified IL-10 protein variants were synthesised by transfecting HEK293T cells with pCCL_wtIL10-tNGFR or pCCL_MQIPQ-IL10-tNGFR (jetPRIME™; VWR) and collecting cell culture supernatants. Control cells were mock transfected. Secreted IL-10 was quantified by LEGENDPlex™ using a 10-plex Human M1/M2 Macrophage Panel.

To test the function of the IL-10 variants, freshly-isolated CD14^+^ cells were cultured overnight in ImmunoCult™ XF supplemented with 100 U/mL penicillin-streptomycin and 5 μg/mL wild-type IL-10 or MQIPQ-modified IL-10. The following day, monocytes were stimulated with 200 ng/mL lipopolysaccharide (LPS) and 5 mM ATP (both from InvivoGen). Monocyte supernatants were collected 5 hours post-stimulation and cytokine production was measured by LEGENDPlex™ using a 10-plex Human M1/M2 Macrophage Panel.

### Suppression assays

All suppression assays were performed in ImmunoCult™ XF supplemented with 100 U/mL penicillin-streptomycin and 5% human serum. For DC suppression assays, Tregs were co-cultured with mature HLA-A2^+^ DCs in the presence of 100 IU/mL IL-2 for 3-days after which the DC phenotype was measured by flow cytometry.

For polyclonal suppression assays, Tregs were labelled with CPD-eFluor™670 (Thermo Fisher Scientific), serially-diluted and co-cultured with allogeneic HLA-A2^−^ PBMCs that were labelled with CPD-eFluor™450 and activated with Dynabeads™ Human T-Expander CD3/CD28 (1:8 bead:PBMC ratio; Thermo Fisher Scientific). Cells were stained and analysed by flow cytometry after 4-days of culture. CD4^+^ Tresponder division was measured using division indices and used to calculate % suppression (inverse of percent Tresponder proliferation) relative to Tresponders cultured alone. Suppression of CD80 and CD86 expression on CD19^+^CD4^−^ B cells in the responder PBMC population was concurrently measured (relative to responders cultured alone).

For antigen-specific alloantigen-dependent suppression assays, HLA-A3^−^ Tregs were labelled with CPD-eFluor™670, serially-diluted and co-cultured with allogeneic HLA-A2^+^ mDCs. After 3-days, CD3^+^ Tresponder cells (previously enriched from an HLA-A3^+^ individual using the EasySep™ Human CD3 Positive Selection Kit II; STEMCELL Technologies) were labelled with CPD-eFluor™450 and added to the co-cultures at a 5:1 Tresponder:DC ratio. Co-cultures were maintained for a further 4-days before being stained and analysed by flow cytometry. % suppression was calculated as described above, gating on HLA-A3^+^CPD-eFluor™670^−^ responder cells.

Antigen-specific GAD65 peptide-dependent suppression assays were performed using frozen CD4^+^ responder T cells that expressed the 4.13-TCR (*30*). To generate these responder T cells, enriched CD4^+^ T cells (EasySep™ Human CD4 Positive Selection Kit II, STEMCELL Technologies) were stimulated with Dynabeads™ Human T-Expander CD3/CD28 (3:1 bead:T cell ratio), lentivirally transduced to express the 4.13-TCR, tNGFR-enriched (Human CD271 MicroBeads Kit, Miltenyi Biotec) and frozen on day 9 of culture (*83*). For suppression assays, Tregs were serially-diluted and co-cultured with HLA-A2^+^HLA-DR4^+^ immature DCs. After 1-2 days, 4.13-TCR^+^ T cells were thawed and added to the co-cultures at a 5:1 Tresponder:DC ratio along with 1 nM GAD65 peptide (NFIRMVISNPAAT). In some experiments, control DCs were pulsed with an irrelevant haemagglutinin (HA) peptide (PKYVKQNTLKLAT). Co-cultures were maintained for a further 2-days before being stained and analysed by flow cytometry. % suppression was determined by measuring inhibition of Ki67 expression in the CD4^+^mTCRβ^+^CPD-eFluor™670^−^ responder T cells.

### *In vivo* A2-CAR Treg chronic stimulation model

To chronically stimulate A2-CAR Tregs *in vivo*, sub-lethally irradiated (150 cGy) A2-NSG mice (10-weeks old) were intravenously (tail vein) administered 1.5-3.2×10^6^ A2-CAR Tregs with an equal number of autologous HLA-A2^−^ PBMCs as an *in vivo* source of IL-2. In these experiments, PD1^KO^ A2-CAR Tregs were not treated with AAV6 thus did not express tCD19. Control mice were administered saline or PBMCs alone. Mice were routinely monitored with no evidence of xenoGVHD development and humanely euthanized 21-days post-infusion. Cardiac blood was collected and red blood cells were lysed with ammonium chloride (STEMCELL Technologies). Spleens were dissociated through 70 µm cell strainers to obtain a single cell suspension. Resulting lymphocytes were stained and analysed by flow cytometry or cell sorted for TSDR analysis.

### TSDR analysis

DNA was isolated and bisulphite converted using the EZ Direct Kit (Zymo Research). The TSDR was PCR-amplified using the AllTaq PCR Core Kit (Qiagen) and the following primers: FWD: AGAAATTTGTGGGGTGGGGTAT), REV (biotinylated): ATCTACATCTAAACCCTATTATCACAACC. PCR products were run on a Q96 MD pyrosequencing system (Qiagen) using the following sequencing primer: AGAAATTTGTGGGGTGGG. Data were analysed using Pyro Q-CpG software (Biotage) by comparing cytosine vs. thymine incorporation at 7 CpG sites in the TSDR.

### Statistical analysis

Flow cytometry data was analysed using FlowJo™ X, LLC (BD Biosciences). Data was analysed using GraphPad Prism 10 (La Jolla) and presented as mean ± SEM with the contribution of each donor shown. Statistical significance was determined using paired two-tailed Student’s t-tests or repeated measures mixed-effects analyses. P-values are provided throughout where p<0.05 was considered significant.

## ACKNOWLEDGEMENTS

This work was supported by grants from Juvenile Diabetes Research Foundation (JDRF) Canada via the JDRF Centre of Excellence at UBC (Grant Key: 3-COE-2022-1103-M-B) and Canadian Institute of Health Research (CIHR; FDN-154304). DAB was supported by fellowships from the Canadian Institute of Health Research (CIHR) and Michael Smith Health Research BC. MKL is Canada Research Chair in Immune Engineering and receives a Scientist Salary Award from the BC Children’s Hospital Research Institute. The study was conceptualised by DAB, JKG, AJL and MKL. Experiments were performed and analysed by DAB, SM, JKG, VCWF, MH, MM, CMW and AB. The manuscript was written by DAB and critically evaluated by MKL. We also thank Lisa Xu for her assistance in cell sorting. MKL has patents pending and a license related to A2-CARs. The other authors have no commercial or financial interests to report.

## SUPPLEMENTAL FIGURES

**Supplemental Figure 1.**
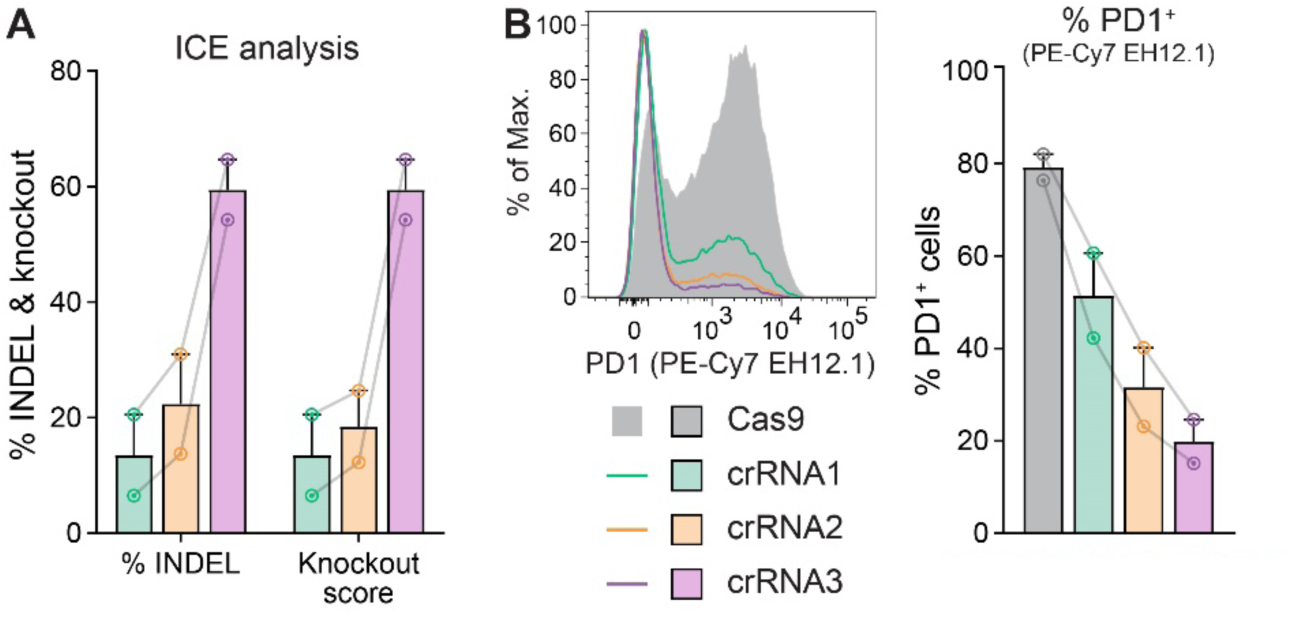
Comparison of gRNAs targeting PDCD1 locus. CD4^+^ T cells were edited with RNPs containing one of three gRNAs targeting the *PDCD1* locus. (**A**) ICE analysis 3 days following electroporation. (**B**) PD1 expression in CD4^+^ T cells 48-hours following polyclonal stimulation. Averaged data are mean + SEM with connected series representing individual subjects (n=2).

**Supplemental Figure 2.**
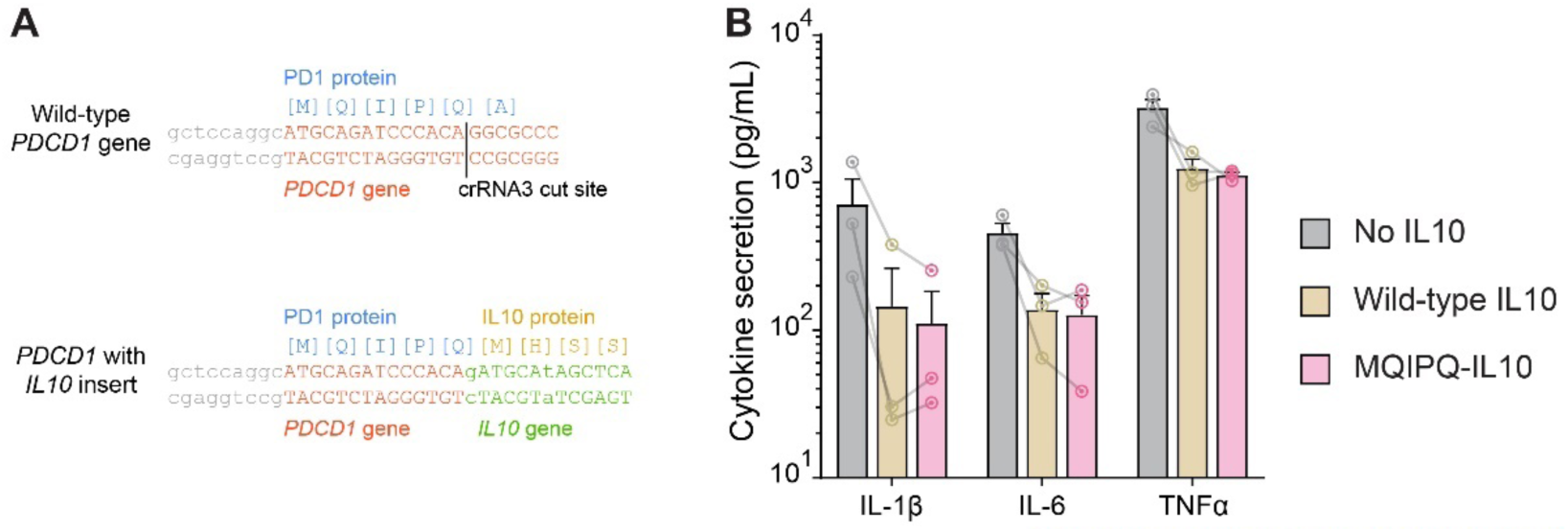
Functional comparison of IL-10 variants. (**A**) Schematic diagram showing the crRNA3 cut site and resulting N-terminal modification of IL-10 when inserted into this locus. (**B**) CD14^+^ monocytes were cultured with 5 µg/mL wild-type or MQIPQ-modified IL-10 overnight and then stimulated with LPS/ATP. Cytokine secretion into cell culture supernatants was measured 5-hours post-stimulation by LEGENDPlex™. Averaged data are mean + SEM with connected series representing individual subjects (n=3).

**Supplemental Figure 3.**
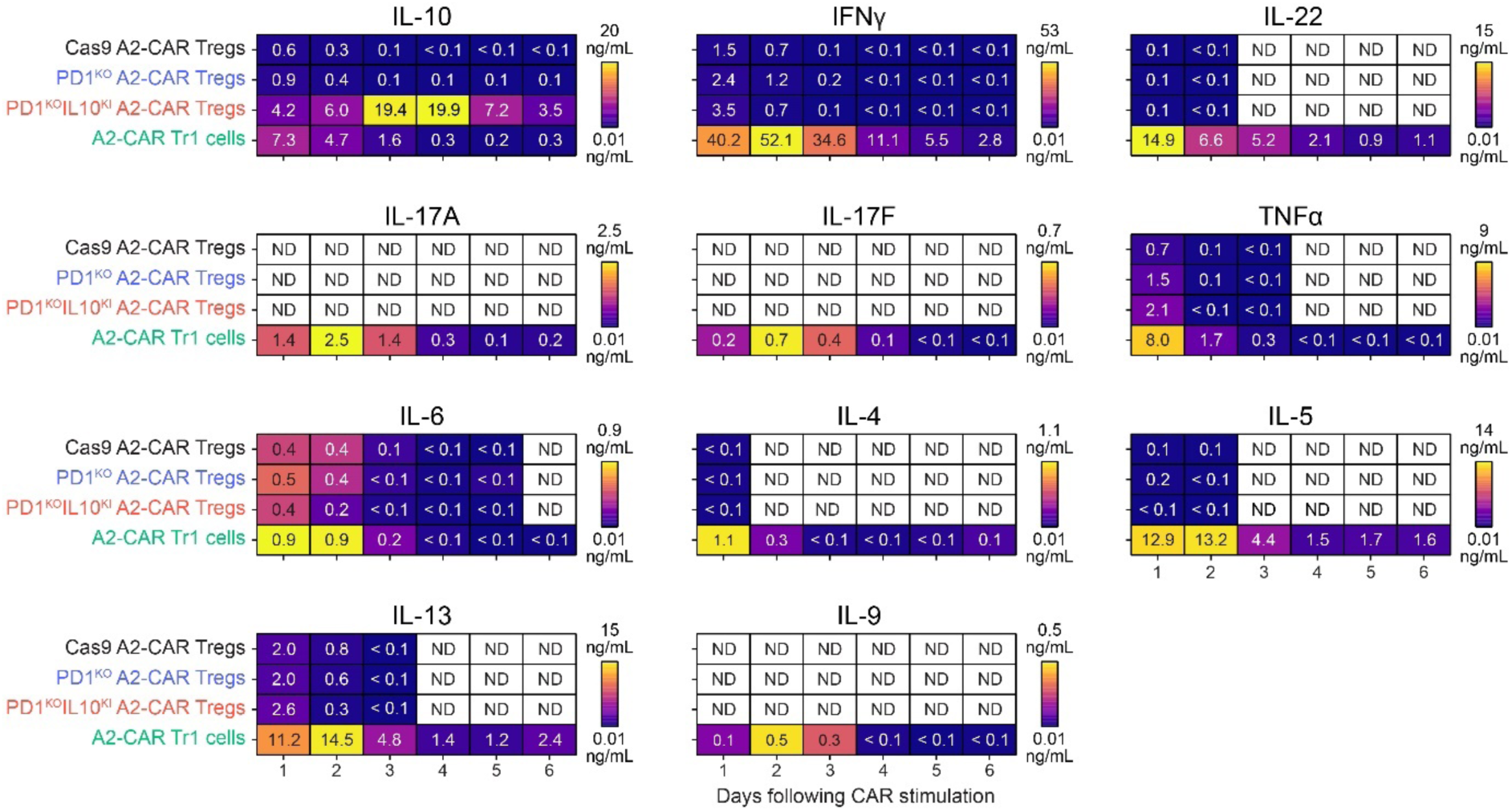
PD1KOIL10KI A2-CAR Tregs secrete minimal proinflammatory cytokines in response to CAR stimulation. A2-CAR Treg and Tr1 cytokine secretion timecourse following co-culture with HLA-A2^+^PD-L1^−^ K562 cells. Every 24-hours, culture supernatants were harvested, the cells were washed twice with PBS and re-cultured in fresh media with IL-2. Data are average of 4 subjects. ND = not detected.

