## Supplemental data for "Fourth generation CAR Tregs with *PDCD1*-driven IL-10 have enhanced suppressive function"

### SUPPLEMENTAL FIGURES

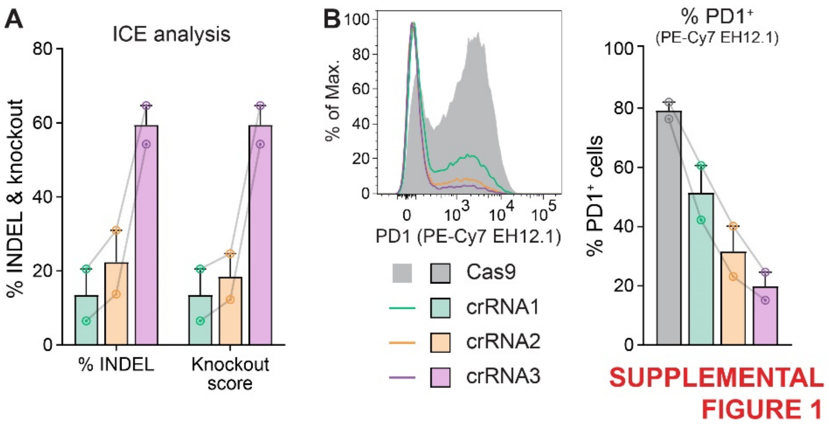


**Supplemental Figure 1** | *Comparison of gRNAs targeting PDCD1 locus*. $CD4^+$ T cells were edited with RNPs containing one of three gRNAs targeting the *PDCD1* locus. (**A**) ICE analysis 3 days following electroporation. (**B**) PD1 expression in $CD4^+$ T cells 48-hours following polyclonal stimulation. Averaged data are mean + SEM with connected series representing individual subjects (n=2).

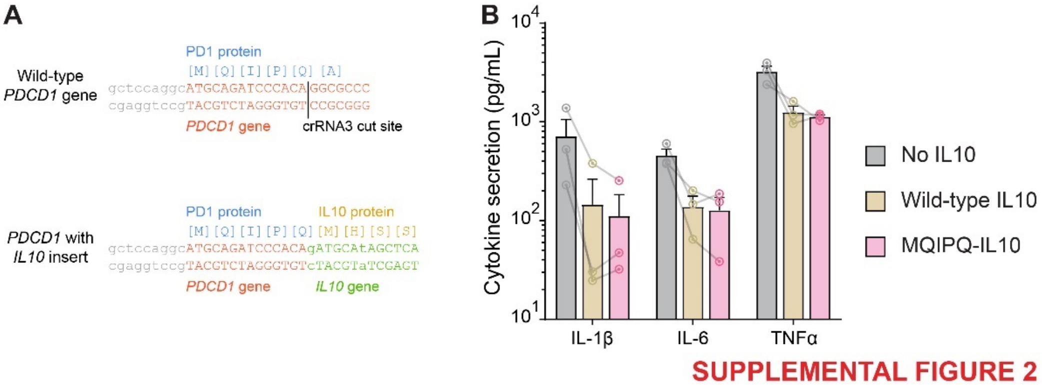


**Supplemental Figure 2** | *Functional comparison of IL-10 variants*. (**A**) Schematic diagram showing the crRNA3 cut site and resulting N-terminal modification of IL-10 when inserted into this locus. (**B**) CD14$^{+}$ monocytes were cultured with 5 µg/mL wild-type or MQIPQ-modified IL-10 overnight and then stimulated with LPS/ATP. Cytokine secretion into cell culture supernatants was measured 5-hours post-stimulation by LEGENDPlex™. Averaged data are mean + SEM with connected series representing individual subjects (n=3).

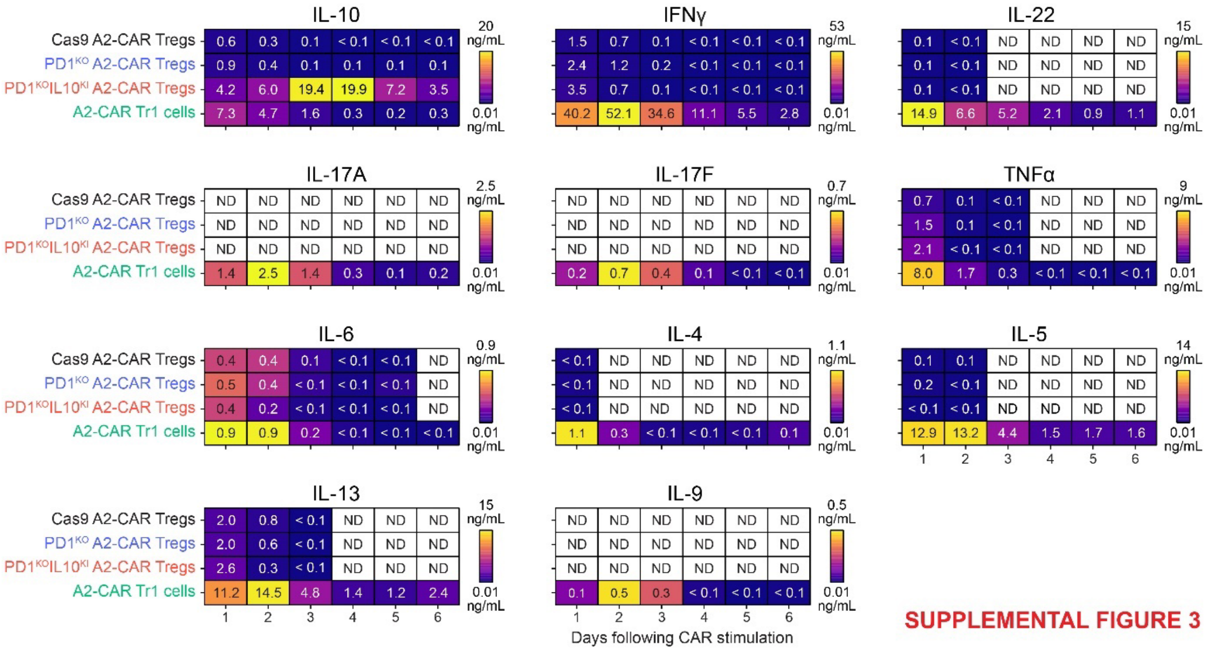


SUPPLEMENTAL FIGURE 3

**Supplemental Figure 3** | *$PD1^{KO}IL10^{KI}$ A2-CAR Tregs secrete minimal proinflammatory cytokines in response to CAR stimulation*. A2-CAR Treg and Tr1 cytokine secretion timecourse following co-culture with HLA-$A2^{+}$PD-$L1^{-}$ K562 cells. Every 24-hours, culture supernatants were harvested, the cells were washed twice with PBS and re-cultured in fresh media with IL-2. Data are average of 4 subjects. ND = not detected.
